# Fatty acid binding protein 1 and fatty acid synthetase overexpression have differential effects on collagen III and cross-linking in Zongdihua pig tissues

**DOI:** 10.1101/2022.06.15.496270

**Authors:** Rong Yang, Di Zhou, Zhihong Yan, Zhonghai Zhao, Yan Wang, Jun Li, Liqun Ren, Lingling Xie, Xin Wang

## Abstract

The purpose of this study was to determine whether FABP1 and FAS regulate expression of collagen and its crosslinking via lysyl oxidase in Zongdihua pigs. We wished to identify processes affecting meat quality using molecular genetics to provide a basis for breeding improvement of these animals. We measured expression levels of FABP1 and related genes using qRT-PCR in longissimus dorsi muscle and subcutaneous adipose tissues. Primary adipocytes from fat tissues were isolated and FABP1 and FAS were overexpressed from recombinant plasmids. Our sequence analysis of the cloned genes indicated that FABP1 gene encodes a hydrophobic protein of 128 amino acids and contained 12 predicted phosphorylation sites and no transmembrane region. FAS encodes 333 amino acid hydrophobic protein containing with 26 phosphorylation sites and 0 transmembrane regions. The basal levels of FABP1 and FAS in pig tissues expression were 3 −3.5-fold higher in subcutaneous fat compared with muscle (P < 0.01). Recombinant expression plasmids were successfully transfected into the cloned preadipocytes and (a) overexpression of FAS resulted in significantly increased expression of COL3A1 gene (P < 0.05) and significantly inhibited lysyl oxidase *LOX* expression (P < 0.01); (b) overexpression of FABP1 significantly increased COL3A1 expression (P < 0.01) and significantly inhibited *LOX* expression (P < 0.05) and significantly reduced lysyl oxidase activity (P < 0.01). Therefore, enhancing FABP1 expression increases collagen accumulation and this preliminarily suggests that FAS and FABP1 can serve as fat-related candidate genes providing a theoretical basis for the study of fat deposition in Zongdihua pigs.

---

The extracellular matrix (ECM) plays an important role in adipocyte proliferation, differentiation and migration and anchors these cells to prevent mechanical movement. The primary component of the ECM is Type III collagen encoded by COL3A1 that is synthesized in large amounts during fat deposition [3]. Lysyl oxidase (LOX) catalyzes collagen and elastin crosslinking and adds to the tensile strength of the ECM [10].The ECM is also a reservoir of components involved in protein-protein interactions such as transforming growth factor β (TGF-β) that is a positive regulator of collagen genes and is highly expressed in tissues with high fat content [4]. These data indicated the presence of an intrinsic link between fat deposition and collagen. In particular, fatty acid binding protein (FABP) family members such as FABP1 play key roles in fat deposition as well as synthesis and degradation of body fat [5–7]. The anabolic enzyme fatty acid synthase (FAS) is also highly expressed in adipocytes and catalyzes triglyceride synthesis and thereby is a regulator of fat deposition [8, 9].

The Zongdihua pig is a local characteristic pig breed of Guizhou, China and possesses the characteristics of a high-grade meat food [11]. In the present study we used this animal as a model of fat deposition on collagen to provide a molecular and genetic basis for improving Zongdihua pig breeds.

## 1 Materials and Methods

### 1.1 Ethics Statemen

This study was carried out in strict accordance with the guidelines for the use of laboratory animals in Guizhou University. This protocol was approved by the Experimental Animal Ethics Review Committee of Guizhou University (protocol number: EAE-GZU-2021-P019). All operations were performed under anesthesia with sodium pentobarbital, and every effort was made to minimize pain.

### 1.2 Test samples

Healthy Zongdi pigs 9-months and 3-days old pigs with similar body weights were obtained from Ziyun County Ziwei Animal Husbandry. Samples (2 g) of subcutaneous fat and longissimus dorsi muscle were collected and placed in an RNA preservation solution (Sartoris. Kibbutz Beit-Harmek, Israel) and then frozen in liquid nitrogen and then stored at −80°C until use.

### 1.3 Reagents

Trizol, Revert Aid First Stand cDNA Reverse Transcription Kit, liposome, mycoplasma remover, PCR TA Cloning Kit, SsoFASt EvaGreen Supermix and 2× Ex Taq Master Mix were purchased from Thermo Scientific (Pittsburg, PA, USA). LOX activity was measured using a commercial kit using the manufacturers protocol (Shanghai Jianglai Biological). PCR amplicons were purified from agarose gels using a purification kit (Vazyme Biotech, Nanjing, China). Ampicillin and kanamycin (Kan) were purchased from Kangwei Century Biotechnology (Beijing, China). Restriction endonucleases *Sal*I and *Kpn*I and T4 DNA ligase were purchased from Qingke Biotechnology (Beijing, China). Fetal bovine serum, Opti-MEM and DMEM/F12 tissue culture media were purchased from Gibco (Grand Island, NY USA). Oil Red O stain was purchased from Solarbio Biotech Company (Beijing, China). Lipofectamine 2000 and 3000 (Invitrogen, Carlsbad, CA, USA) were used for recombinant plasmid transfections.

### 1.4 Primer design and synthesis

Primer 5 software (http://www.premierbiosoft.com/primerdesign/) was used to design primers for the porcine GAPDH (XM_021091114.1) FABP1 and FAS genes from the respective GenBank sequences NM_001004046.2 and NM_213839.1. GAPDH served as internal reference gene (XM_021091114.1). All primers were synthesized by Qingke Biotechnology (Table 1).

**Table 1.**
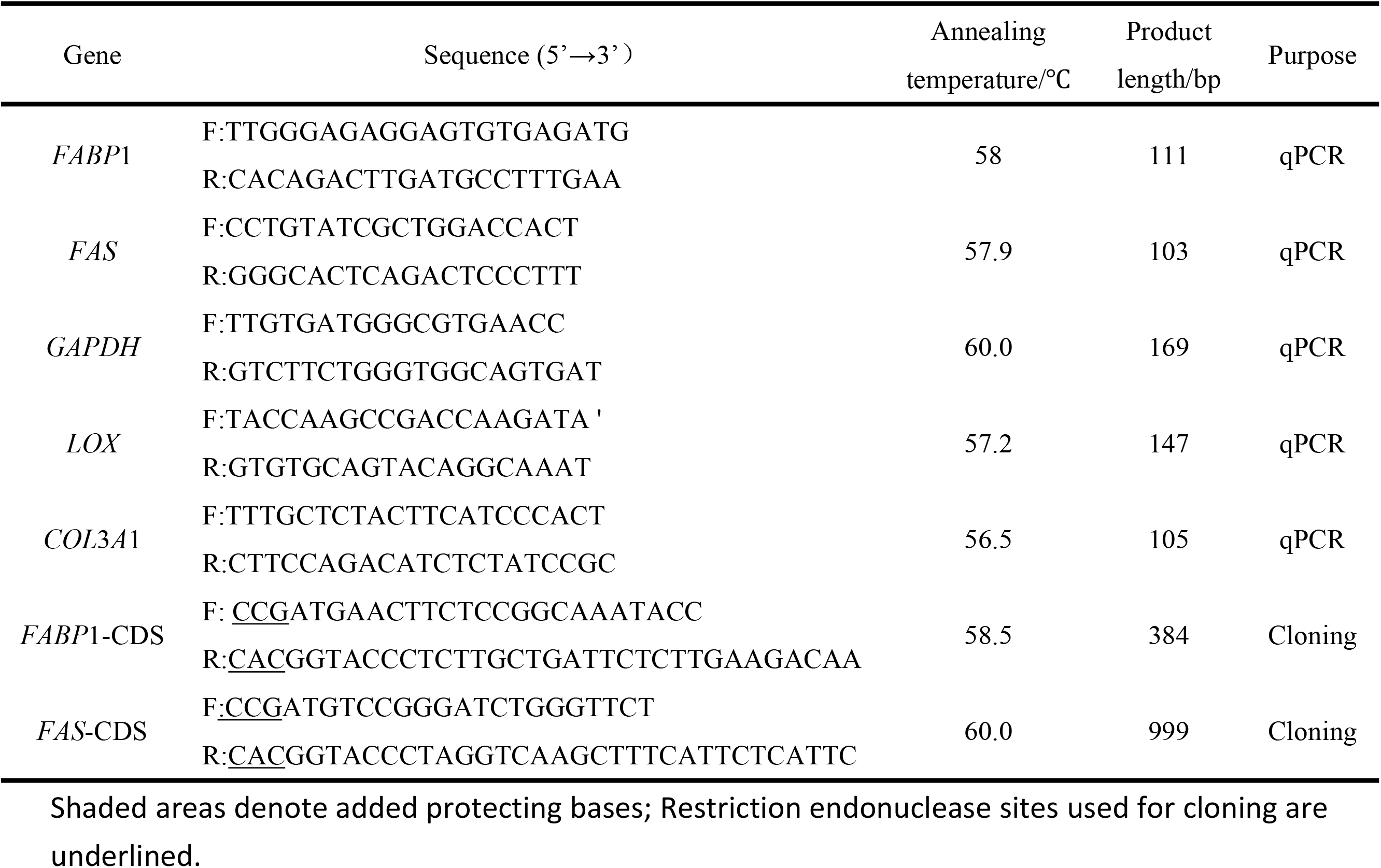
PCR primers used in this study.

### 1.5 RNA extraction and mRNA expression analysis

Total RNA was extracted from homogenized fat and longissimus dorsi muscle using Trizol and the protocol recommended by the manufacturer (Invitrogen, Carlsbad, CA, USA). First strand cDNA was synthesized using the HiFi Script cDNA kit as recommended by the manufacturer (Thermo Fisher). The concentration and purity of RNA and cDNA was determined using a micro-UV spectrophotometer (Nanodrop, Thermo Fisher). Real-time quantitative PCR (qRT-PCR) was used to measure steady state mRNA levels from tissues and cells (see below) with GAPDH as the internal reference gene and FABP1 and FAS as the quantitative targets. The reactions (10 μL) in triplicate contained 5 μL SsoFASt EvaGreen Supermix, 5 pmol each primer, and 1 μL cDNA template that were cycled at 95°C for 3 min and 39 cycles of 95°C for 15 s, 60°C for 15 s and 72°C for 1 min and collection time for 5 sec. Melting curve analysis to determine amplicon purity was performed at increments of 0.5°C from 60 to 95°C. The 2-^ΔΔ^Ct method was used to process and analyze the data.

### 1.6 Construction of FABP1 and FAS plasmid expression vectors

Subcutaneous fat cDNA was used as template to PCR-amplify the CDS regions of the FABP1 and FAS genes. Amplicons were generated using the 2× Ex Taq kit (Thermo) with the following conditions: 95°C for 5 min and 35 cycles of 95°C for 30 s, 60°C for 45 s and 72°C for 30 s followed by a final extension at 72°C for 5 min. The PCR products were recovered from agarose gels using a commercial kit (see above) and cloned using the TA vector pUCM-T vector (Bio Basic, Ontario, Canada) and transformed into TOP 10 competent cells (Invitrogen) using the manufacturers specifications. The transformation mixes were plated on ampicillin agar plates and incubated at 37 °C overnight. Single colonies were incubated in Luria Bertani broth containing ampicillin at 37°C for 8 h. Possession of cloned genes were verified by PCR and further sequenced by Qingke Biological to confirm insert identity. The cloned inserts were released by PstI and KpnI digestion and cloned into the eukaryotic expression vector pEGFP-C1 (Clontech) using the above procedure but with kanamycin selection. Plasmid inserts were verified by sequencing as per above.

### 1.7 Isolation and culture of adipocytes

Healthy 3-day-old Zongdihua pigs were used for isolation of adipose cells. Subcutaneous adipose tissues were collected and quickly washed in PBS lacking Ca^2+^ and Mg^2+^ and connective tissues and blood vessels were removed. This tissue was then cut into pieces and digested with type II collagenase for 90 min. The digestion was terminated by adding DMEM/F 12 medium containing 10% serum and the mixture was filtered through sterile gauze and a cell sieve (200-mesh and 400-mesh). Cells from the filtrate were collected by centrifugation at 1500 rpm for 10 min and suspended in serum-free medium, mixed by pipetting and then centrifuged as per above. The cell pellets were suspended in 10% DMEM / F12 complete medium and transferred to a cell culture flask and placed at 37°C in a 5 % CO_2_ atmosphere for 4 h. The medium was replaced and the cells were incubated as per above. When the cells reached approximately 85% confluence the cells were removed using trypsin / EDTA and transferred to 6-well plates and grown to 90 % confluence. The medium was replaced and culture was continued for a total of 2-3 days. Differentiation was induced by replacing the growth media with induction medium (10% DMEM/F12 complete medium, DEX stock solution, IBMX concentrated stock solution and insulin stock solution) and changed every 2 days to induce differentiation. The cells were observed for the production of lipid droplets using an inverted microscope. When lipid droplets were present in about 80 % of the cells, Oil Red O staining was performed to identify adipocytes. The amount of stain present in the wells was quantified using spectrometry at 510 nm in a microplate reader.

### 1.8 Transfection of cultured adipocytes

Cultures identified as adipocytes were transfected at 80% confluence by replacing the maintenance medium with the following components: recombinant plasmids pEGFP-C1-FABP1 and pEGFP-C1-FAS separately were combined with 125 μL OPTI-MEM, 5 μL P3000 and 4 μg DNA. Separately, in 5 μL Lipofectamine 2000 125 μL OPTI-MEM were combined. The mixtures were then combined and mixed and placed at 37°C in a 5% CO_2_ atmosphere for 15 min. Transfected cells were inoculated in 6-well plates that was cultures as per above for 24 h and examined for LOX activity (see below).

The culture medium from transfected cells were used to assay for LOX enzyme activity using a commercial kit (see above) and activity was measured using absorbance at 450 nm in a microplate reader. The concentration of porcine lysyl oxidase (LOX) activity in the sample was calculated according to a standard curve constructed using Excel

### 1.9 qRT-PCR

Transfected cells (see above) were used to assess the effects of the overexpression of FABP1 and FAS on COL3A1 and LOX gene expression. Cells were plated in 6-well plates and to grown to 90% confluence and transfected as per above. The cells were removed after 36 h at 37°C and steady state mRNA was measured using qRT-PCR(Section 1.4).

### 1.10 Data processing and analysis

The 2^−ΔΔ^ Ct method was used to determine relative mRNA expression levels of FABP1 and FAS. SPSS20.0 software (IBM, Chicago, Ill, USA) was used to determining significance levels. P < 0.01 indicated very significant differences and P < 0.05 significant differences. DNAStar (Madison, WI, USA) was used for DNA sequence analysis.

## 2 Results and Analysis

### 2.1 Differential expression analysis

Our preliminary experiments examined the expression of FABP1, FAS, LOX and COL3A1 in subcutaneous fat and longissimus dorsi muscle tissues of the Zongdihua pigs. The expression levels for all 4 genes were extremely significantly (P < 0.01) elevated in subcutaneous fat tissues compared with the muscle (control) tissues (Figure 1). We therefore generated PCR amplicons for FABP1 and FAS to be used for cloning and over-expression analysis. We successfully amplified FAS and FABP1 (Figure 2) and they were then cloned using the TA vector pUCM-T. The genes were then transferred to the pEGFP-C1 expression vector and inserts were verified by sequence analysis and the cloned inserts were free of any mutations when compared with the reference sequences. The predicted FAS gene encoded a protein of 333 amino acids (37.593 kDa) with pI =7.377, pH=7.0. The FABP1 gene product was composed of 128 amino acids (14.107 kDa) and pI = 7.094(Figure 3). FAS contained 22 potential S/T phosphorylation sites and 4 potential tyrosine phosphorylation sites while FABP1 possessed 12 S/T phosphorylation sites (Figure 4). Additional analyses of predicted structures of the FAS and FABP1 proteins indicated they were typical for these proteins in other pig breeds.

**Figure 1.**
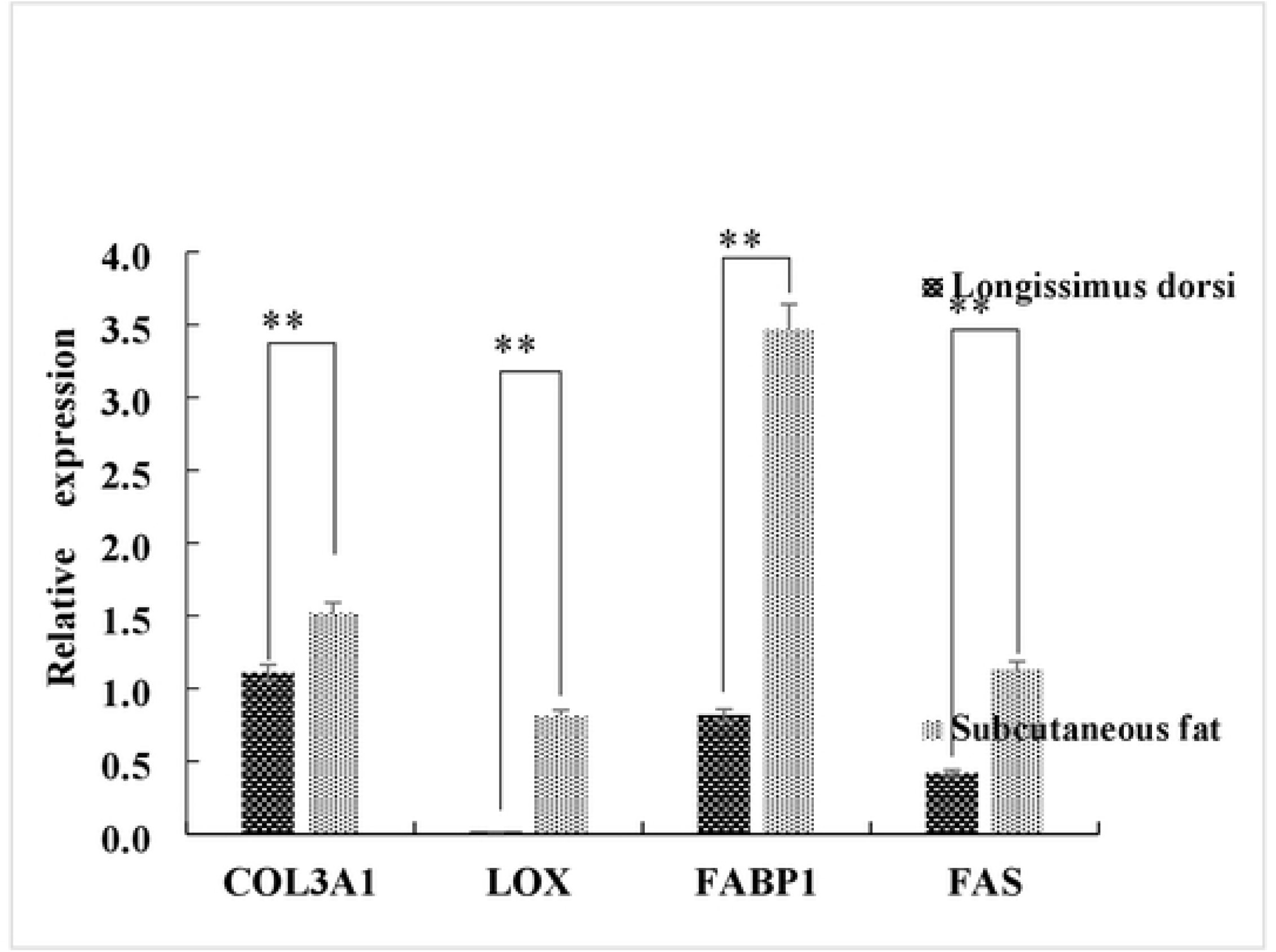
Fat deposition and relative expression of collagen-related genes in porcine adipose and muscle tissue. **, p<0.01.

**Figure 2.**
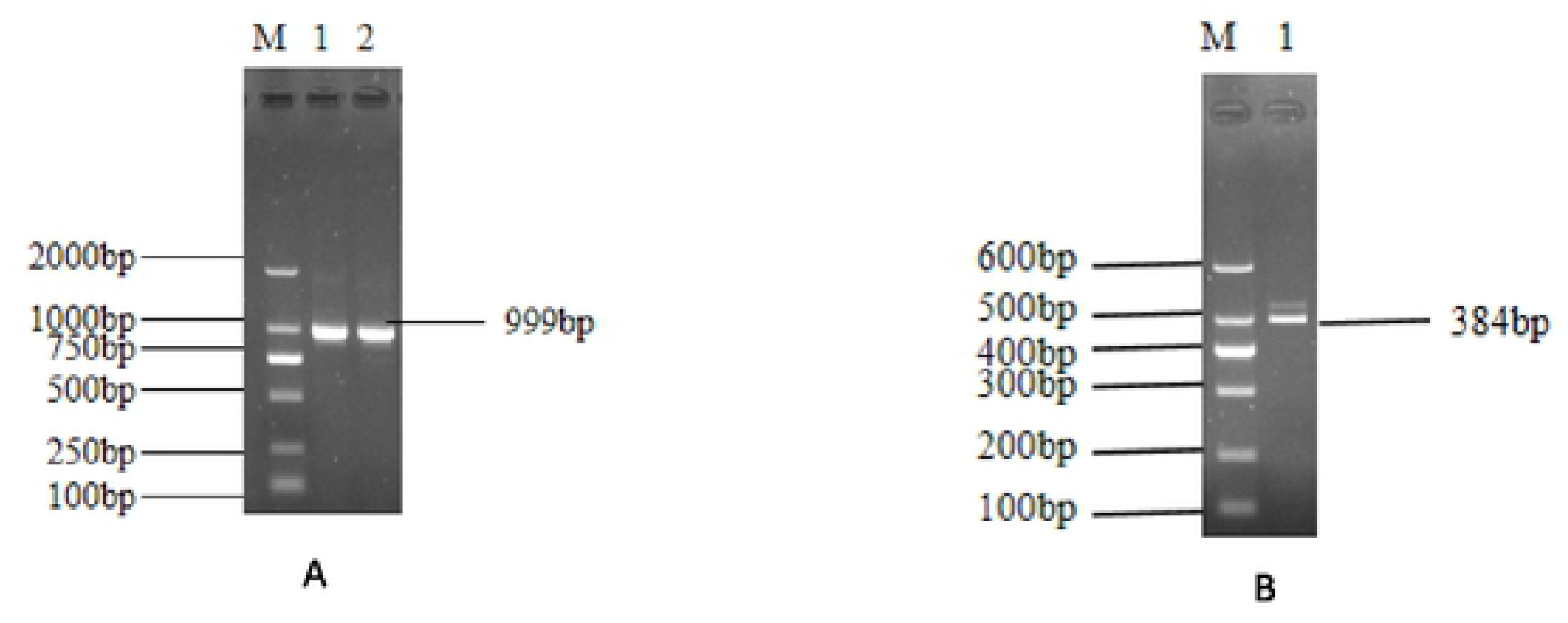
PCR amplification of (A) *FAS* and (B) *FABP*1. Amplicon and molecular size standards are indicated in bp.

**Figure 3.**
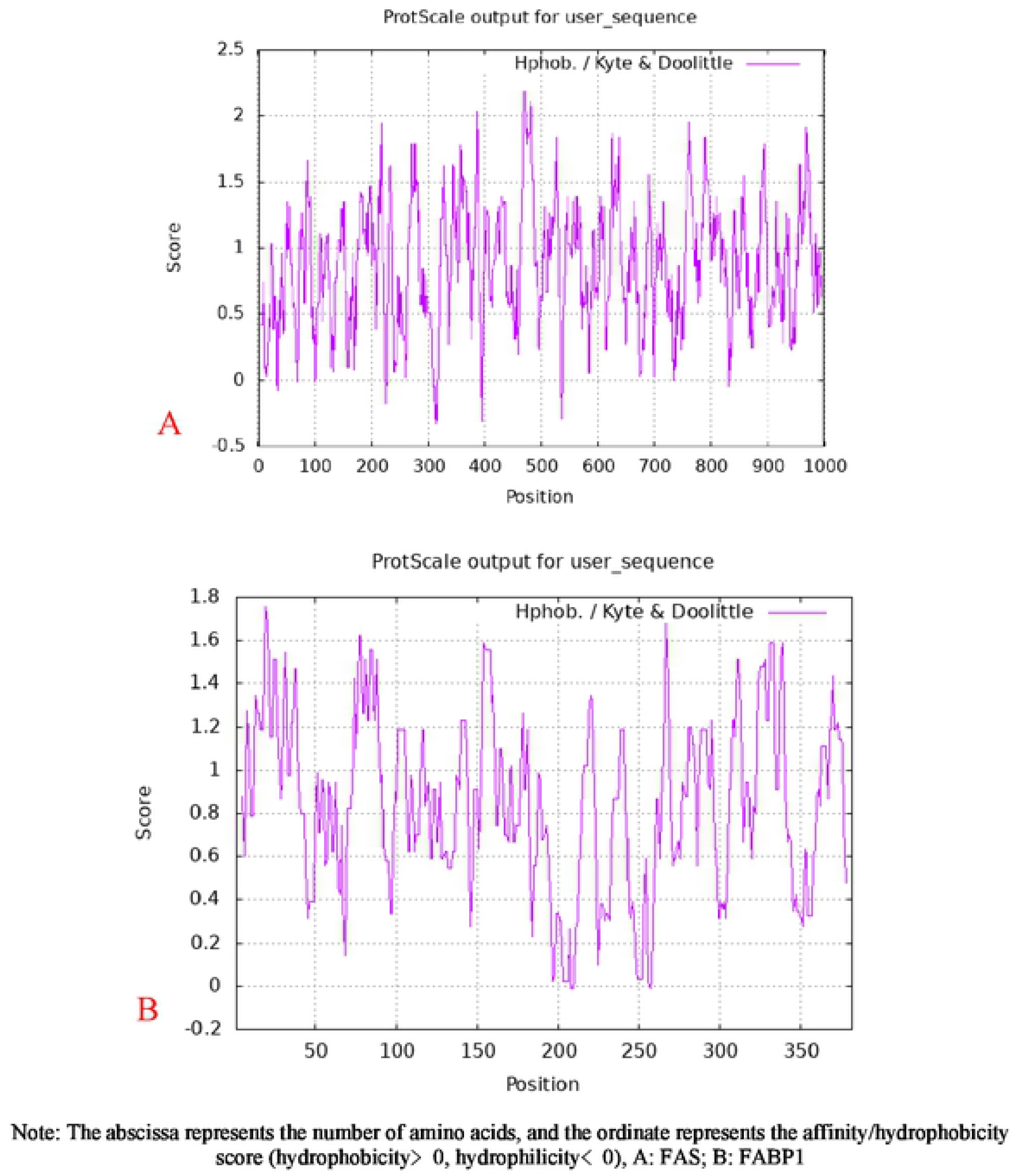
The affinity/hydrophobicity analysis of FAS and FABP1 proteins

**Figure 4.**
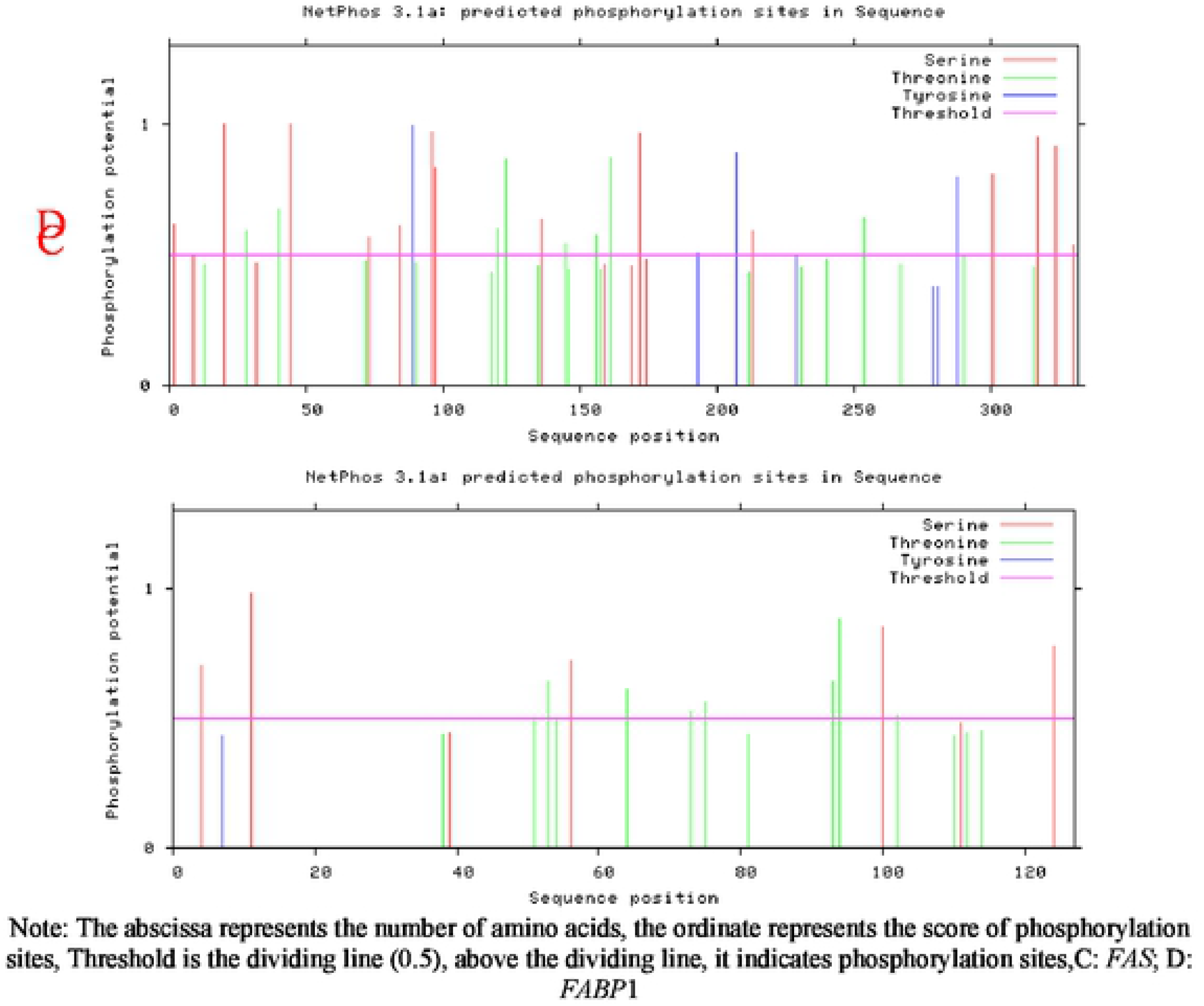
Analysis of FAS and FABP1 protein phosphorylation sites

### 2.2 Adipocyte culture and analysis

We utilized the subcutaneous fat of Zongdihua pigs to isolate pure cultures of adipocytes using differential adhesion. The subcutaneous adipocytes appeared round and adhered to the cell culture flask after 4 h in culture and were 80 % confluent after 24 h. A subset of the cells appeared as long spindles with transparent bodies (Figure 5A and 3B). After 48 h in culture, the cells had proliferated and most of the cells appeared as long spindles and these reached confluence by 72 h (Figure 5C and 3D).

**Figure 5.**
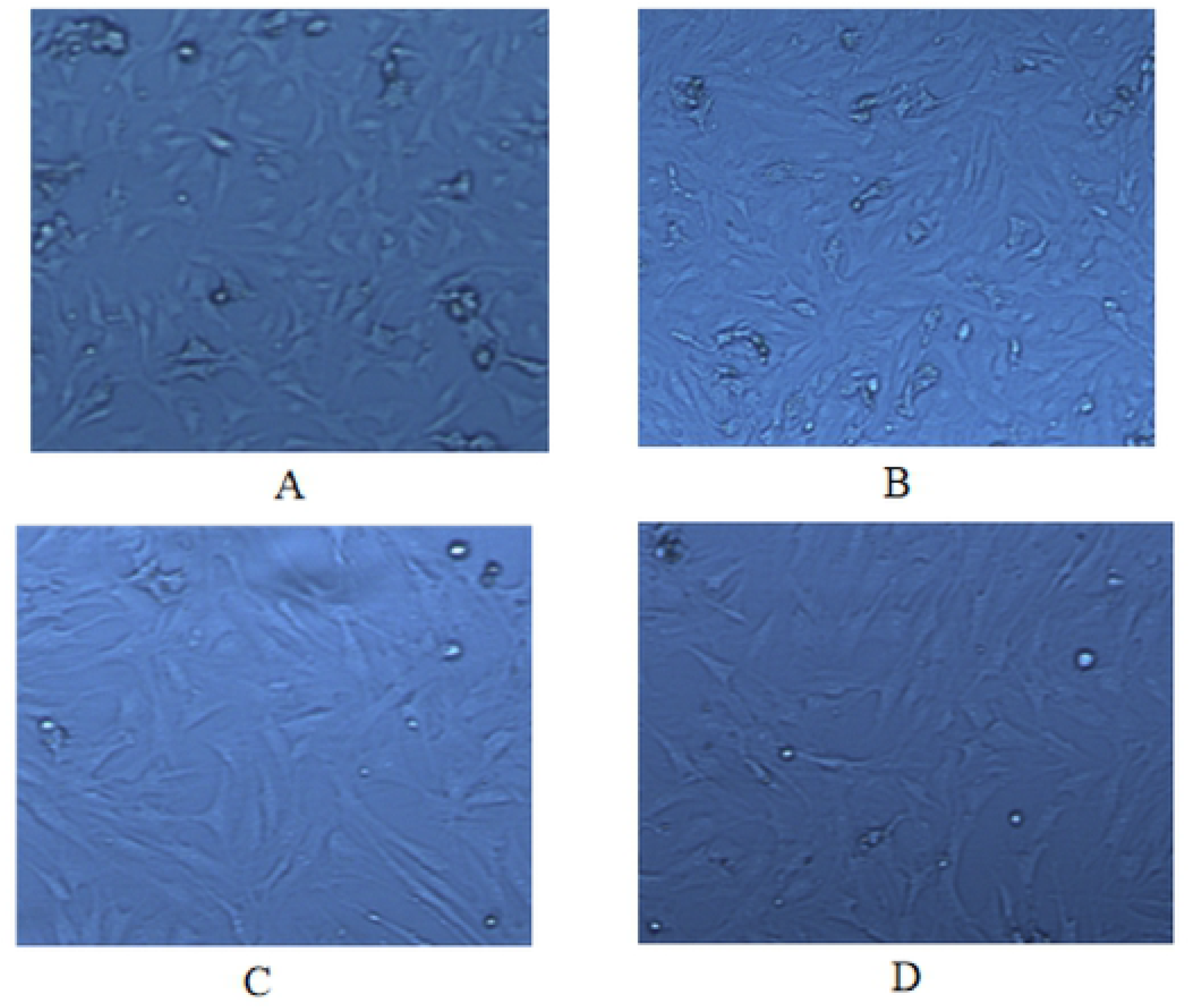
Morphological changes of pre-adipocyte primary cultures. Photomicrographs of cultures taken after (A) 4 h (B) 24 h (C) 48 h and (D)72 h. 100× magnification.

The identity of these cells as adipocyte precursors could be confirmed if they were able to differentiate in the presence of differentiation induction medium, stain with Oil red O and differentiate. We found that the long spindles of the precursor cells gradually became elliptical following 24 h exposure to the differentiation medium (Figure 6A). Lipid droplets began to appear in the cells 72-96 h (Figure 6B). After 6 days of induction and culture, the number of lipid droplets increased significantly and the volume increased (Figure 6C). The lipid droplets fused with the cells after 8 d. At this time, differentiation maintenance medium was replaced with induction medium and the cultures were allowed to continue for another 4 - 5 d. The round and transparent bodies in the cells were verified as lipid droplets by oil red O staining (Figure 6D). This confirmed that our primary cultures were differentiated into adipocytes.

**Figure 6.**
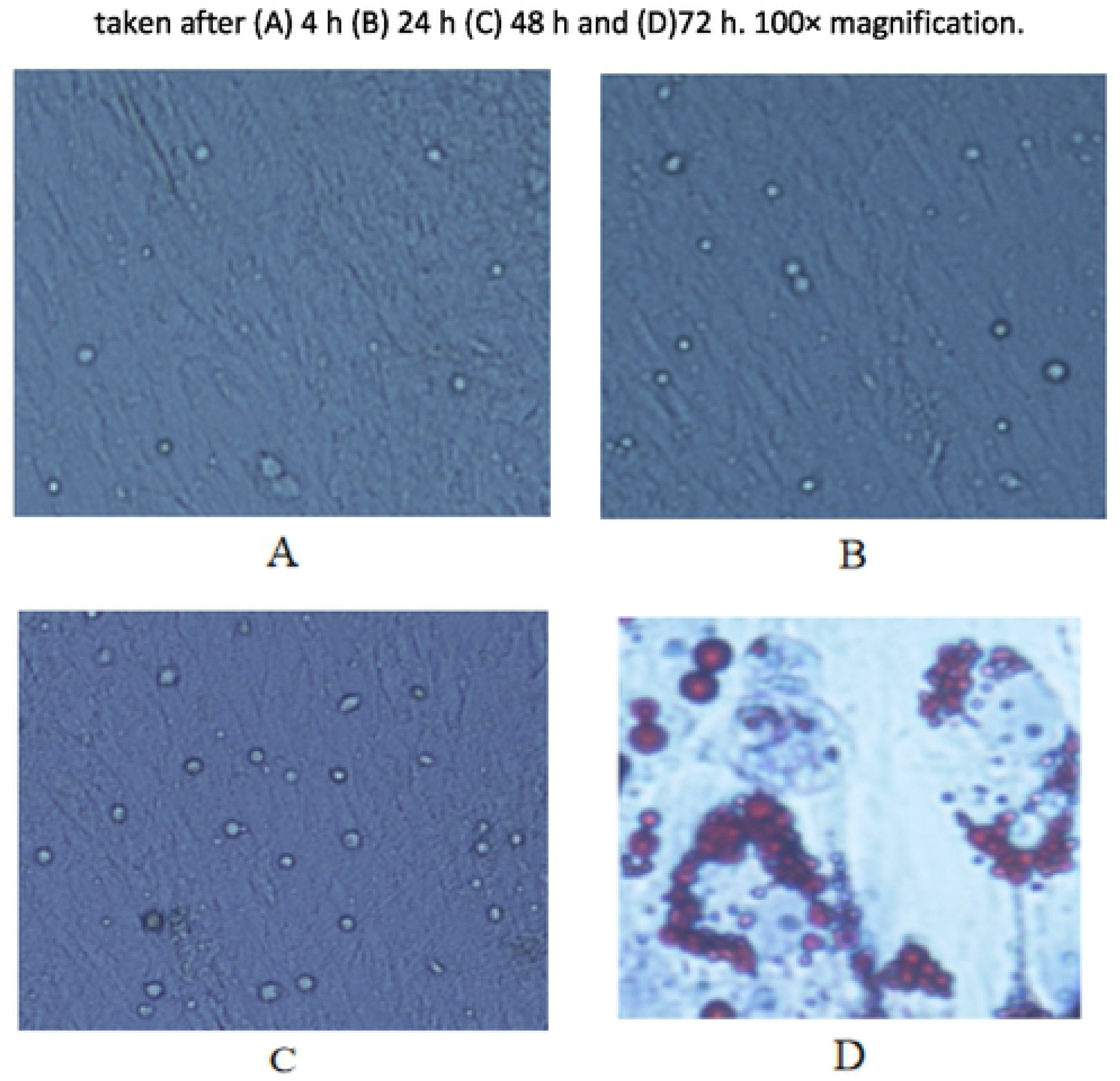
Induced differentiation of intramuscular preadipocytes. Photomicrographs of differentiating cells at (A) 24 h (B) 4 d and (C) 6 d. (D) Oil red O staining of a cell monolayer after 5d in differentiation medium. 100 × magnification.

### 2.3 Transient over-expression analysis of FAS and FABP1 in differentiated adipocytes

Cloned versions of FAS and FABP1 were constructed that were fused to the eGFP protein at their carboxyl termini and over-expressed via the strong CMV promoter. The transfection procedure for the cells was followed using fluorescent microscopy and 24h following transfection, clear and bright green protein expression was seen for both plasmid constructs compared with cells transfected with the vector only or with a blank control (Figure 7A-5E). This indicated that the eGFP fusions were successfully expressed in the cells.

**Figure 7.**
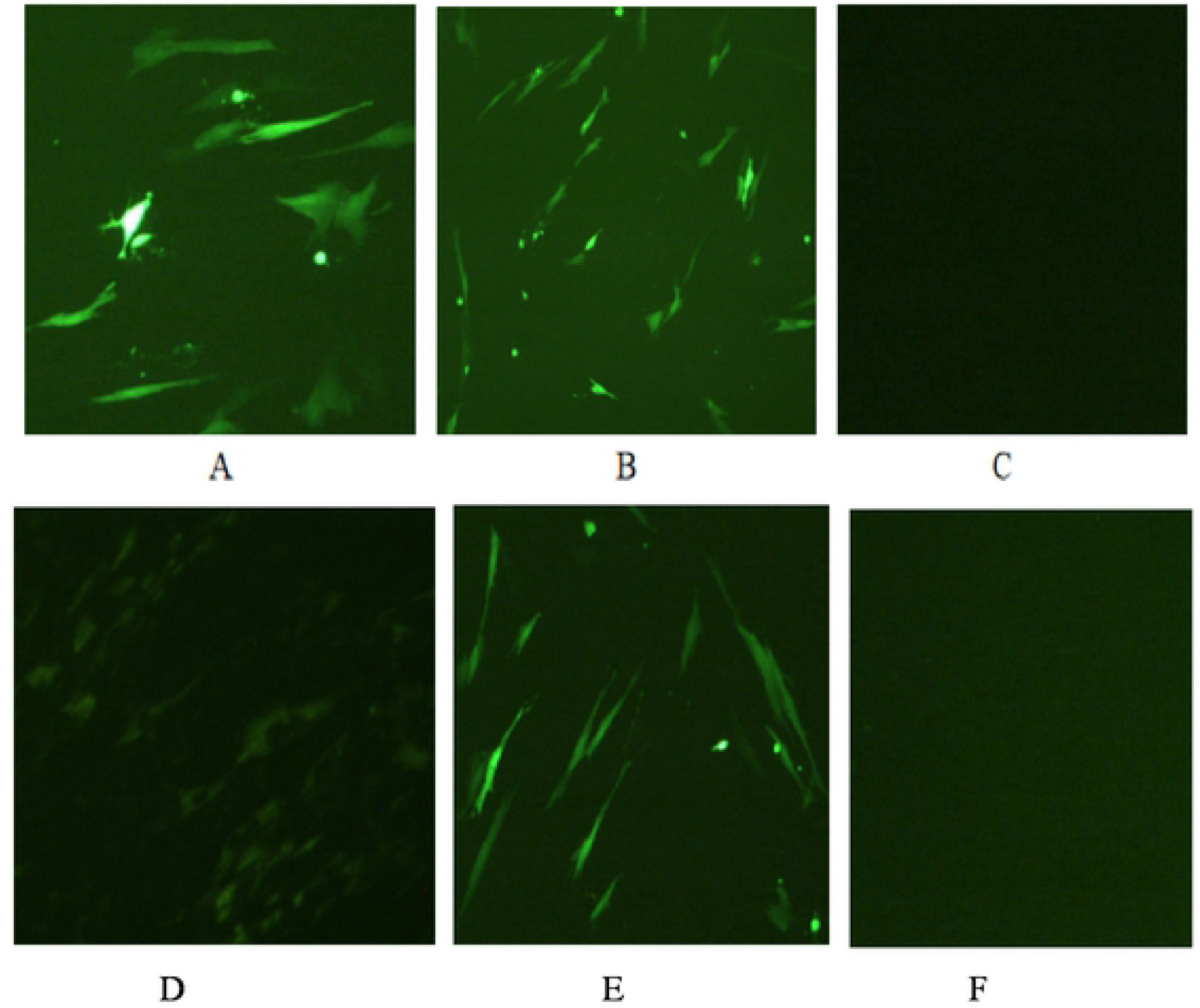
(A) pEGFP-C1-*FAS* transfected cells (B) vector only (C) blank control (D) pEGFP-C1-*FAS* (E) vector only (F) blank control.

These cells were then examined for the ability of the cloned genes to stimulate expression of *COL3A1* and *LOX* using real-time quantitative PCR. We found that 36 h following transfection, mRNA levels for both *FAS* and *FABP1* were 3 - 3.5-fold higher in the transfected cells versus controls (P<0.01). Interestingly, over-expression of the cloned genes resulted in significantly higher levels for *COL3A1* (P < 0.05) and lower levels for *LOX* (P<0.01) (Figure 8).

**Figure 8.**
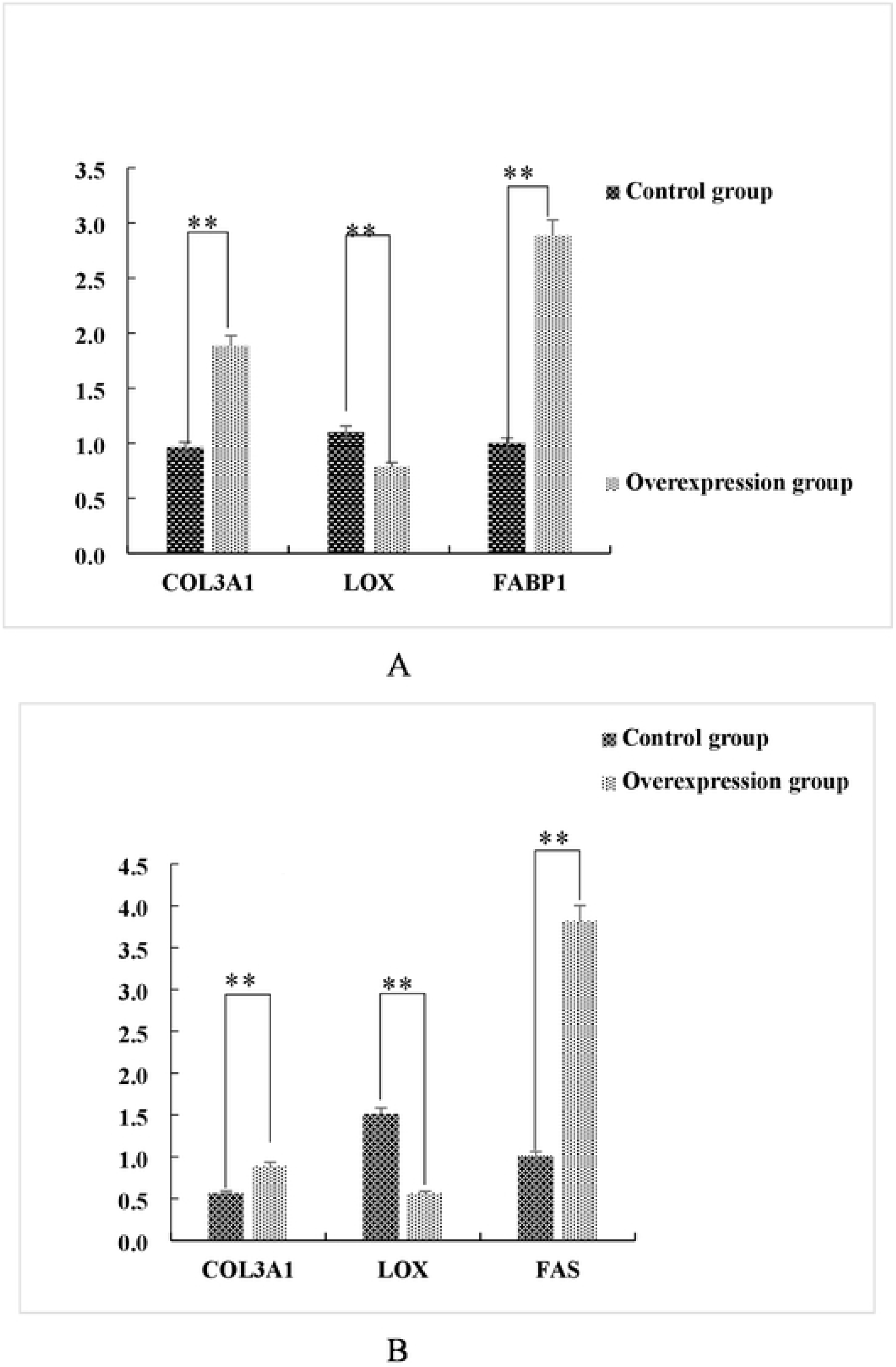
Over-expression of FAS and FABP1 stimulate Col3A1 and inhibit LOX expression. Cultured adipocytes were transiently transfected with pEGFP-C1-FAS and pEGFP-C1-FAS as indicated(A and B). Steady state mRNA levels for the indicated genes were determined using real time qRT-PCR. *, p < 0.05; **, *p < 0.01*.

We further examined whether the effects of the transfected plasmids on the decreases in *LOX* mRNA levels were also reflected in decreased lysyl oxidase activity. The supernatants of the transfected cells used above were examined for lysyl oxidase activity using a double-antibody sandwich method. We generated a linear standard curve for the assay (Figure 9A). Interestingly, the lysyl oxidase activity mirrored those of the mRNA levels and LOX enzyme activity decreased to levels that were significantly (P < 0.01) lower for the transfectants compared with controls (Figure 9B and 9C).

**Figure 9.**
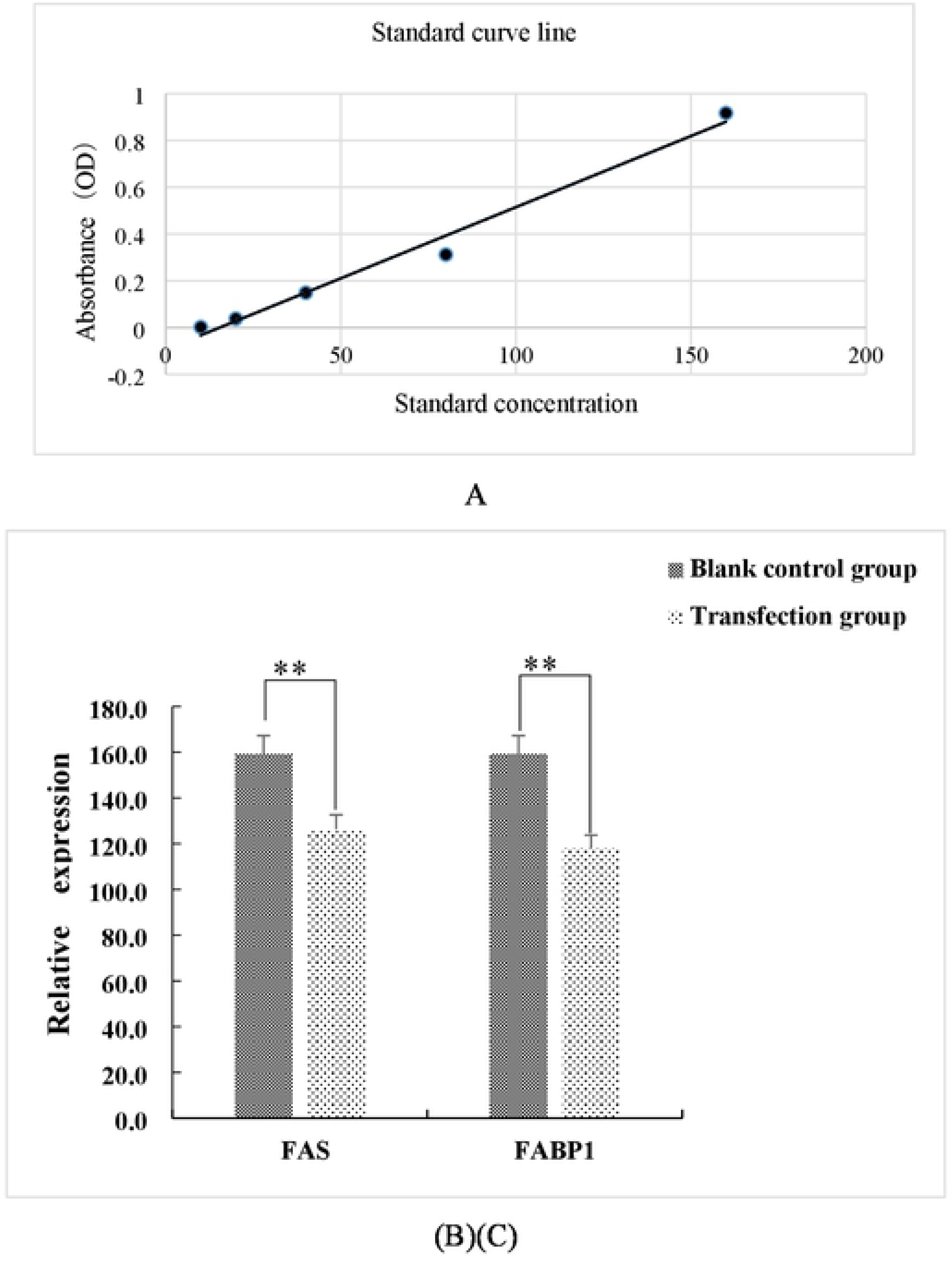
Lysyl oxidase activity is decreased in cultured adipocytes over-expression either FAS of FABP1 (A). Standard curve for lysyl oxidase activity. LOX activity measured in cells transfected with (B) FAS and (C) FABP1. **, *p < 0.01*.

## 3 Discussion

The FABP gene family encodes fatty acid-binding proteins that are involved in the transport of intracellular fatty acids as well as the synthesis and degradation of fat in the body. Therefore, this family plays an important role in the fat metabolism in the body [12]. FABP1 plays an important regulatory role in fat deposition and participates in the formation of triglycerides [13]. Elevated expression of FABP1 in porcine preadipocytes significantly increases fat accumulation [14]. In goats, the FABP1 gene in subcutaneous fat is also expressed at significantly higher levels than in the control longissimus dorsi muscle (P<0.05) [15]. FAS is a key enzyme in fat synthesis in animals and promotes fatty acid formation. The FAS gene is expressed in mammalian fat, kidney and lungs and other tissues and cells [16]. FAS is also highly expressed in the subcutaneous adipose tissue of Baixi and first generation Subai hybrid pigs [17] in addition to Bamei and large white breeds [18–19]. In this study, we found that FABP1 and FAS gene expression were significantly higher than for the control muscle tissue (P<0.01). This indicated that FABP1 and FAS could be used as candidate genes for fat deposition studies especially since higher expression levels correlated with greater fat deposition. Our *in-silico* bioinformatic analysis of FABP1 and FAS proteins indicated slight differences but the overall structures were similar to their counterparts in other pig breeds. We therefore used these subcutaneous fat tissues to culture primary adipocytes to enable more directed and comprehensive experimentation. Our results agreed with previous studies indicating that COL3A1 and LOC gene expression in subcutaneous fat are significantly higher and lower than in longissimus dorsi muscle tissues, respectively. These results are consistent with the higher levels of collagen crosslinking in muscle compared with fat tissues. Therefore, LOX activity and fat deposition are negatively correlated and fat deposition may damage the structure of connective tissues resulting in an improvement in tenderness with age [23]. Therefore, fat deposition can alter collagen cross-linking but further research is required to determined direct causality.

We were able to successfully over-express FAS and FABP1 in Zongdihua pig preadipocytes and resulted in increases in *COL3A1* and decreases in *LOX* expression (P<0.01) as well as lysyl oxidase activity. Therefore, fat deposition may promote *COL3A1* gene expression thereby promoting collagen production and inhibiting LOX gene expression. Overexpression of FAS gene can promote cell proliferation, inhibit apoptosis and elevate triglyceride synthesis in adiposities. These gene products thereby collectively affect the deposition and decomposition of fat [24]. Collagen is an ECM protein that not only supports adipocytes, but also regulates the development of adipose tissue via cell-cell signaling [25]. Disruption of collagen III synthesis may lead to impaired triglyceride accumulation in adipocytes and disrupt adipose tissue remodeling [26]. A previous study has demonstrated a role for adipose tissue in regulating insulin sensitivity that can also enhance the beneficial metabolic effects of branched-chain fatty acid esters of hydroxy fatty acids, and therefore enhance anti-inflammatory effects [27]. Lysyl oxidase (LOX) and 4 other lysyl oxidase-like proteins (LOXL 1-4) are copper amine oxidases with a highly conserved catalytic domain that use lysine tyrosyl quinone cofactors and possess conserved copper binding sites. This group also catalyzes the first step in covalent cross-linking of ECM proteins such as collagen and elastin that are responsible for mechanical strength of the ECM [28. 29]. Our results indicated that overexpression of FAS and FABP1 genes significantly increased the expression of *COL3A1* gene while significantly inhibiting *LOX* gene. This leads to the conclusion that cellular FAS and FABP1 levels can regulate collagen accumulation. However, fat deposition can also promote collagen synthesis but this is negatively correlated with the expression of lysyl oxidase activity and mRNA accumulation. This indicates that fat deposition has an inhibitory effect on LOX and decreased levels of crosslinking would result in a weaker collagen network structure. This is therefore also related to the fat of Zongdihua pigs that results in a thick and waxy skin and enhanced meat tenderness.

## 4 Conclusions

The current study demonstrated that the expression of FAS and FABP1 genes was higher in subcutaneous fat of Zongzi pigs compared with muscle tissue. Overexpression of FAS and FABP1 significantly increased expression of *COL3A1* and significantly inhibited LOX expression and its corresponding lysyl oxidase activity. Our findings suggest that the FAS and FABP1 genes may be fat-related candidate gene targets.

